# APRIL: Adaptive Regression-Based Two-Dimensional Quantitative Anisotropy Imaging Using Acoustic Radiation Force Impulse

**DOI:** 10.64898/2026.06.30.735710

**Authors:** Md Walid Hassan, Kayla Crook, Young Jin Gi, Jangsoon Lee, Md Murad Hossain

## Abstract

**Objective:** This study aims to develop and validate a quantitative, depth-resolved anisotropy imaging framework that extends ARFI-based focal degree-of-anisotropy (DoA) estimation into two-dimensional mapping by modeling the depth-dependent relationship between shear modulus ratio (SMR) and peak displacement ratio (PDR).

**Methods:** We propose APRIL (<u>A</u>daptive <u>P</u>olynomial <u>R</u>egression for anisotropy Imaging via ARFI-induced Disp<u>L</u>acements), a framework for quantitative, depth-resolved DoA imaging that adaptively selects polynomial regression or shape-preserving spline interpolation based on excitation PSF asymmetry. Training data were generated using an LS-DYNA3D + Field II simulation pipeline in homogeneous transversely isotropic media (SMR 0.9–4.9). Testing included shifted SMRs under varied acoustic conditions and three heterogeneous inclusion configurations (anisotropic inclusion in isotropic background and vice versa). Experimental validation was performed in an in-vivo murine tumor model over the time, ex-vivo chicken breast, and tissue-mimicking gelatin phantoms, using a Verasonics system with an L11-5v transducer.

**Results:** APRIL achieved depth-resolved SMR prediction errors below 9% over 10–30 mm, with highest accuracy in the focal region (MAE 2.3%, RMSE *<* 0.1) and stable performance across PSF transition zones. In heterogeneous phantoms, it reconstructed anisotropy maps with SSIM up to 86% and MPE below 7%, accurately delineating inclusion boundaries. Under acoustic parameter variations, mean absolute errors remained below 10%, demonstrating robustness to system and tissue heterogeneity.

**Conclusion:** APRIL enables robust, two-dimensional anisotropy imaging beyond focal estimates.

**Significance:** The method provides a physically grounded and generalizable framework for clinically viable anisotropy biomarkers in muscle, tendon, kidney, tumor and breast tissues.

**Highlights:**

- Novelty: APRIL introduces LoA-conditioned adaptive polynomial-spline regression to extend ARFI-based anisotropy estimation from focal point estimates into full 2D depth-resolved SMR imaging.
- Results: APRIL achieved SMR prediction errors below 9% over 10–30 mm, SSIM up to 86% in heterogeneous phantoms, MAE below 10% under acoustic variations, tracked tumor anisotropy progression in vivo, and differentiated anisotropic inclusion versus isotropic background in tissue-mimicking gelatin phantom.
- Significance: APRIL enables clinically viable, spatially resolved anisotropy biomarker imaging in muscle, tendon, kidney, and tumor tissues without requiring heterogeneous training data.

## I. Introduction

Many tissues—including muscle [1], tendon [2], kidney [3], myocardium [4], brain white matter [5], bone [6], and breast tumors [7]—exhibit anisotropy, with mechanical properties differing between longitudinal (structure-aligned) and transverse (perpendicular) directions due to organized microarchitecture. In each of these cases, directional dependence of stiffness has diagnostic implications. For example, malignant breast tumors are more anisotropic than benign masses [8], [9]. Skeletal muscle anisotropy is disrupted in boys with Duchenne muscular dystrophy, reflecting microstructural degeneration [10]. In kidney cortex, anisotropy is altered by fibrosis, blood pressure, and ischemia-reperfusion injury [3]. In myocardium, anisotropy reflects fiber remodeling in hypertrophic cardiomyopathy [4]. These findings underscore the importance of not only measuring stiffness but also quantifying anisotropy as a biomarker. Mechanical properties of biological tissues reflect underlying microstructural organization and are altered by pathological processes such as fibrosis, tumor infiltration, and ischemic injury, making quantitative mechanical characterization a valuable biomarker for disease diagnosis and monitoring [11]. Acoustic radiation force (ARF)–based ultrasound elastography methods primarily fall into two categories: shear wave elastography (SWE), which estimates tissue stiffness by tracking laterally propagating shear waves induced by ARF, and acoustic radiation force impulse (ARFI) imaging, which evaluates localized, on-axis displacement responses at the excitation site. While SWE provides quantitative shear wave velocity measurements over broader regions, it relies on lateral wave propagation and is susceptible to attenuation, reflections, and directional bias; in contrast, ARFI interrogates tissue mechanics directly along the beam axis with higher spatial resolution and reduced dependence on shear wave tracking. In ultrasound, SWEI has been widely used to assess tissue DoA by measuring shear wave velocities along longitudinal and transverse directions [1]. The methodological considerations and practical limitations associated with lateral shear wave tracking in anisotropic media have been discussed extensively in prior SWEI-based DoA studies [12]–[14], and are therefore not reiterated here.

Acoustic radiation force impulse (ARFI) imaging interrogates tissue mechanics using on-axis displacement responses to the excitation beam [15], [16], reducing dependence on lateral shear wave propagation and improving spatial resolution. In anisotropic tissues, ARFI-derived degree of anisotropy has been quantified using the peak displacement ratio (PDR) obtained from excitations applied parallel and perpendicular to the axis of symmetry. Prior studies modeled tissue behavior using a transversely isotropic (TI) material framework, in which longitudinal and transverse shear moduli (*µ*_*L*_, *µ*_*T*_) define the shear modulus ratio (SMR = *µ*_*L*_*/µ*_*T*_), and demonstrated that PDR is monotonically related to SMR under TI assumptions [17]. Empirical polynomial mappings between PDR and SMR were shown to enable quantitative SMR estimation in swine muscle and kidney [18]. However, both ARFI- and SWE-based approaches have thus far provided focal or point-wise SMR estimates rather than spatially resolved anisotropy images. Generating two-dimensional SMR maps would allow visualization of anisotropic inclusions within isotropic backgrounds and vice versa, thereby extending TI-based modeling from parameter estimation to quantitative anisotropy imaging. However, no existing ARFI-based framework has extended focal anisotropy estimation into spatially resolved two-dimensional SMR imaging conditioned on depth-varying PSF geometry. Prior approaches assume a uniform PDR–SMR polynomial relationship across depth, which breaks down in geometric transition regions where PSF asymmetry reverses. This gap of the absence of a depth-aware, spatially resolved anisotropy imaging framework grounded in ARFI physics-motivates the present work.

In ARFI-based anisotropy estimation, the excitation point spread function (PSF) plays a critical role in shaping the measured peak displacement ratio (PDR). Even for a fixed shear modulus ratio (SMR), variations in PSF geometry across depth alter the relative distribution of acoustic radiation force in the lateral and elevational dimensions, thereby modifying the observed displacement response. In prior work, it is demonstrated under a transversely isotropic (TI) material framework that the ARFI-derived PDR is monotonically related to SMR when PSF geometry is held constant [17]. However, because the excitation beam undergoes geometric convergence and divergence around the focal zone, the relative dominance of the elevational and lateral beamwidths changes with depth. We quantify this geometric asymmetry using the level of asymmetry (LoA), defined as the ratio of elevational to lateral −6 dB beamwidths of the ARF excitation PSF. LoA *>* 1 indicates elevational dominance, LoA *<* 1 indicates lateral dominance, and LoA *≈*1 corresponds to near-symmetric excitation. As LoA varies, the functional relationship between PDR and SMR shifts, even when intrinsic tissue anisotropy remains unchanged. Therefore, accurate depth-resolved SMR imaging requires conditioning the PDR–SMR mapping on PSF asymmetry.

To generate an SMR image from PDR, the relationship between SMR and PDR must be determined at each depth, since the PSF varies with depth. Because of this variation, the relationship is not uniformly a third-order polynomial; even when it follows a third-order polynomial at different depths, the coefficients differ (see Fig. 2). Therefore, robust regression models must be empirically derived at each depth to construct an accurate SMR image. Towards this goal, we propose APRIL: <u>A</u>daptive <u>P</u>olynomial <u>R</u>egression for anisotropy Imaging via ARFI-induced Disp<u>L</u>acements in this study. APRIL uses the level of asymmetry (LoA) of the ARFI PSF and PDR profiles along depth to predict SMR along depth. By learning from simulated TI phantoms spanning a broad range of SMR values, focal depths, SNR, and acoustic parameters, APRIL is trained to predict SMR along the depth axis. APRIL is evaluated on simulated homogeneous and heterogeneous data, and experimentally validated in in-vivo murine tumor models, tissue-mimicking gelatin phantoms, and Sx-vivo chicken breast. Note that APRIL is trained exclusively on homogeneous tissue under fixed nominal acoustic conditions, but tested on both homogeneous and heterogeneous configurations and under varying acoustic parameters, demonstrating generalization to unseen tissue structures and acoustic conditions without requiring heterogeneous or acoustically varied training data.

## II. Methodology

### A. APRIL Framework

The proposed APRIL framework is summarized in Fig. 1. APRIL builds LoA-conditioned mappings between ARFI-PDR and SMR over axial depth, using a two-stage strategy:

**Fig. 1.**
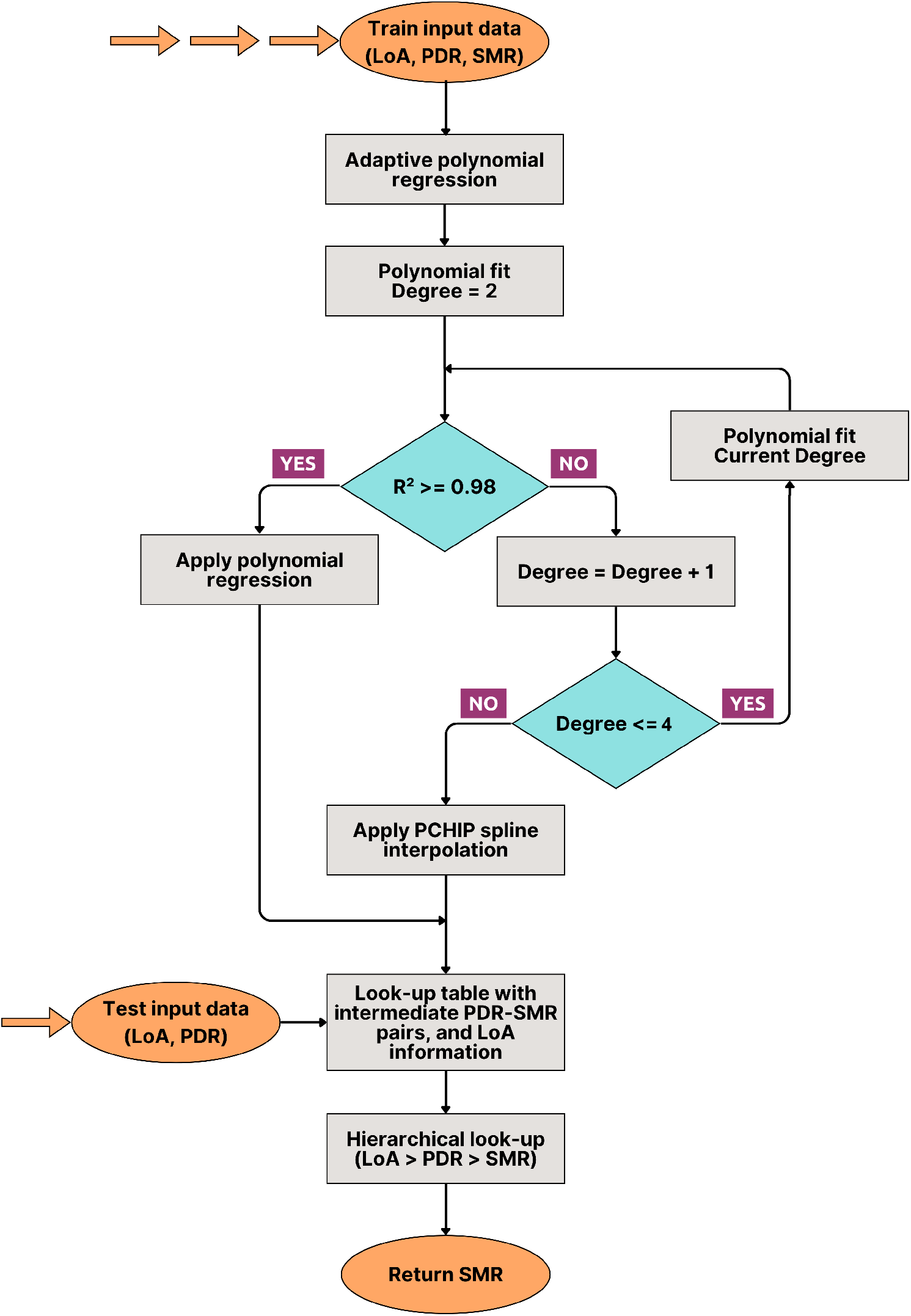
Overall framework of the proposed APRIL method.

#### 1) Adaptive polynomial regression

For each LoA condition, separate polynomial models of increasing degree (upto 4) are fit to the available PDR–SMR pairs. The lowest-degree model that exceeds a predetermined coefficient of determination (R^2^ ≤0.98) threshold is selected for that LoA condition. This high threshold in R^2^ ensures polynomial regression reflects the correct relationship between PDR and SMR.

#### 2) Spline fallback

If polynomial regression fails to meet the (R^2^) requirement or exhibits oscillatory behavior (See Fig. 2 bottom panel), a shape-preserving Piecewise Cubic Hermite Interpolating Polynomial (PCHIP) is used as a fallback. This ensures monotonicity and stability without extrapolation beyond the support of the training data. Because beam focusing produces a stable and smoothly varying radiation force distribution at the focal depth, the PDR–SMR relationship follows a consistent polynomial trend there, whereas away from focus—particularly in geometric transition regions—PSF asymmetry and beam divergence alter force distribution, causing deviations from polynomial behavior.

**Fig. 2.**
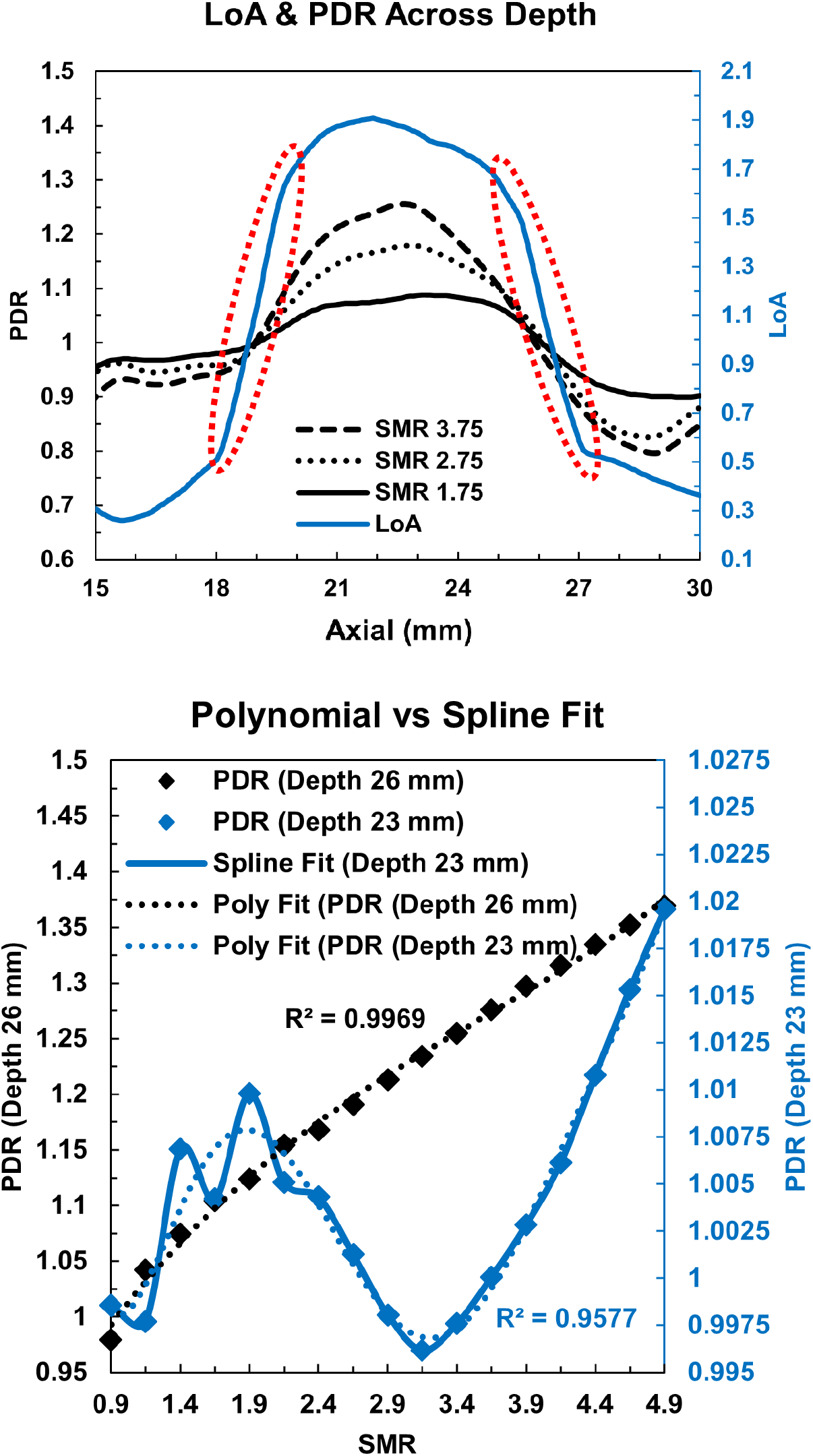
(Top Panel) LoA and PDR across depth from 15 mm to 30 mm for focus 26 mm, SMR 1.75, 2.75, & 3.75, with speed of sound 1540 m/s^**2**^, attenuation coefficient 0.5 dB/cm/MHz, and density 1000 kg/m^**3**^. The dotted elliptical annotations indicate the geometrical transition region of PSF. (Bottom Panel) PDR versus SMR fit at focal depth 26 mm for two representative axial depths: 26 mm (focal region-polynomial 3rd order fit) and 23 mm (PSF transition region-spline fit).

The geometric reversal produces a sharp transition region (see Fig. 2 top panel) along depth where the relative dominance of the lateral and elevational dimensions alternates. Within this zone, the PDR–SMR relationship departs from smooth polynomial behavior, rendering polynomial regression unstable or inaccurate. For these cases, PCHIP interpolation was applied. Each fitted curve (polynomial or PCHIP) was stored together with metadata, including LoA, focal depth, and a densely sampled set of PDR–SMR pairs from the polynomial fit or spline interpolation. At inference, APRIL selects the nearest LoA-conditioned model, and performs a closest-match lookup between the observed PDR values and the precomputed dense PDR–SMR curves, thereby estimating PDR at any depth.

This design provides a computationally efficient, depth-resolved SMR estimation pipeline that is robust to PSF geometry variations across the imaging depth. By precomputing and storing LoA-conditioned PDR–SMR mappings, APRIL avoids repeated model fitting during inference and instead performs rapid curve selection and lookup-based estimation at each axial location. This makes the framework suitable for generating continuous SMR profiles and two-dimensional anisotropy maps with low computational overhead. The combination of LoA-conditioned curve selection and domain-adaptive transformation ensures consistent SMR estimation performance across varying transducer geometries, focal configurations, and imaging depths without requiring retraining for every new test case. Moreover, by explicitly incorporating PSF asymmetry into the estimation process, the method reduces sensitivity to depth-dependent beam-shape transitions that would otherwise introduce bias in conventional single-polynomial PDR–SMR models. As a result, APRIL provides a stable and physically interpretable pathway for extending focal ARFI-based anisotropy estimation into spatially resolved quantitative anisotropy imaging.

### B. Dataset Preparation

Datasets were generated using finite-element method simulations of ARFI excitation in transversely isotropic (TI) phantoms modeled using LS-DYNA3D (Livermore Software Technology Corp. Livermore, CA), and Field II [19] [20] pipeline and ultrasound tracking, following established models of impulsive acoustic radiation force in anisotropic elastic media [15], [17], [18].

The excitation and track frequencies are 4 MHz and 6 MHz with F-numbers 1.5 and 1.25, respectively. The excitation pulse duration was 70 *µ*s. The span of the FEM mesh in axial, lateral, and elevational directions was 2 mm to 42 mm, −6 mm to 6 mm, and −5 mm to 5 mm with an isotropic element size of 0.2 × 0.2 × 0.2 mm^3^.

The LS-DYNA3D `MAT_ORTHOTROPIC_ELASTIC` material model was used to simulate TI elastic materials. Only homogeneous materials were used for training with SMR: 0.9–4.9, step 0.25. The speed of sound, attenuation, and density were fixed to 1540 m/s, 0.5 dB/cm/MHz, and 1000 kg/m^3^, respectively, for the train materials. The focal depth was varied to 18 and 30 mm. APRIL was tested on both homogeneous and heterogeneous materials with focal depth at 22 and 25 mm, respectively. Homogeneous materials had SMR of 1.25-4.75, step 0.5 with variable speed of sound (1500, 1540, 1580 m/s), attenuation (0.25, 0.5, 1 dB/cm/MHz), and density (900, 1000, 1100 kg/m^3^). Three heterogeneous phantoms were simulated with inclusion’s SMR of 1, 2, and 4 embedded in a background with SMR of 4, 1, and 1, respectively. For all materials, the longitudinal-to-transverse Young’s modulus ratio was fixed at *E*_*L*_*/E*_*T*_ = 1.01 (*E*_*L*_ = 12.8 kPa, *E*_*T*_ = 12.2 kPa), and the transverse shear modulus was fixed at *µ*_*T*_ = 4 kPa. The Poisson’s ratio in the plane of symmetry was set to *ν*_*LT*_ = 0.499, the Poisson’s ratio in the plane of isotropy was calculated as 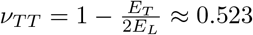, and the reciprocal ratio was given by 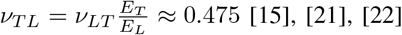.

The level of asymmetry (LoA) was calculated as the ratio of the elevational to lateral −6 dB beamwidths of the ARF excitation point spread function (PSF) at each depth, where LoA *>* 1 indicates elevational dominance, LoA *<* 1 indicates lateral dominance, and LoA = 1 corresponds to near-symmetric PSFs. Displacement data were analyzed over an axial range spanning 10–30 mm for test evaluation, with LoA computed across the same depths. Note that FEM mesh displacement output extends to 32 mm; however, the evaluation range was restricted to 10–30 mm to exclude near-field artifacts and regions of negligible acoustic energy at the distal boundary. The framework then used these inputs to generate continuous, depth-resolved shear modulus ratio (SMR) estimates, or aggregated into 2D anisotropy maps.

To evaluate SMR, the ARFI PSF was oriented both parallel and perpendicular to the material AoS. The resulting displacement fields from the FEM simulations were then used to drive scatterer motion in Field II, where ultrasound radiofrequency (RF) data were synthesized using the L7-4 imaging parameters [16] mentioned previously. Each Field II simulation employed a density of 25 scatterers per resolution cell, which were generated with distinct random scatterer distributions. To approximate realistic acquisition conditions, additive white Gaussian noise (AWGN) was applied to every RF line using MATLAB (MathWorks, Natick, MA), resulting in a system echo signal-to-noise ratio (SNR) of 25 dB, which is consistent with typical clinical ultrasound systems and human tissue.

Axial motion tracking was performed using one-dimensional normalized cross-correlation as described in [16]. This yielded a two-dimensional (axial × time) dataset of displacement waveforms for each simulated transversely isotropic (TI) material and each PSF orientation. From these displacement profiles, the peak displacement (PD) was identified.

### C. Simulated Test Configurations

Simulation-based evaluation was performed using Verasonics Vantage system (Verasonics Inc., Kirkland, WA) equipped with L7-4 transducer parameters under three test scenarios:

1. Homogeneous validation: SMR values of 1.25:0.5:4.25 were tested at focal depth 22 mm with fixed acoustic properties (speed of sound 1540 m/s, density 1000 kg/m^3^, attenuation 0.5 dB/cm/MHz).
2. Acoustic parameter variation: For SMR values 1.75, 2.75, and 3.75, robustness was evaluated under varying speed of sound (1500, 1540, 1580 m/s), density (900, 1000, 1100 kg/m^3^), and attenuation (0.25, 0.5, 1 dB/cm/MHz).
3. Heterogeneous phantoms: Two cases of anisotropic inclusions (SMR = 2 and 4) embedded in isotropic background (SMR = 1), and one case of isotropic inclusion (SMR = 1) embedded in anisotropic background (SMR = 4), were evaluated at depth 25 mm.

### D. Experimental Setup

All experimental acquisitions were performed using a Verasonics Vantage ultrasound research system (Verasonics Inc., Kirkland, WA) equipped with an L11-5v linear array transducer. The transducer was mounted on a Velmex rotary table (Velmex Inc., Bloomfield, NY) integrated with a Velmex BiSlide three-axis mechanical translational stage, enabling precise and repeatable angular and translational positioning. The complete motion control system was mounted on a vibration isolation table (Newport Corp., Irvine, CA) and controlled programmatically via MATLAB (MathWorks, Natick, MA). ARFI push and tracking frequencies were set to 4.75 MHz and 7.6 MHz, respectively. ARFI excitations were applied at multiple angular orientations relative to the dominant tissue axis, and PDR was computed from the maximum and minimum peak displacement responses across all acquired orientations, approximating the parallel and perpendicular responses without requiring prior knowledge of the tissue axis of symmetry. To test the data acquired with L11-5v transducer, we simulated training data only for focal depth 18 and 25mm with L11-5v transducer parameters.

#### 1) Ex Vivo Chicken Breast

Fresh chicken breast specimens were obtained commercially and imaged within 24 hours of purchase. Each specimen was positioned in a water bath at room temperature (~22°C) with the dominant fiber orientation identified visually from surface striations. Chicken breast pectoralis major muscle exhibits transverse isotropy with longitudinal shear modulus *G*_*t*_ = 33.59 *±* 14.13 kPa and transverse shear modulus *G*_*p*_ = 13.52 *±* 0.91 kPa (mean*±* SE, *n* = 3) measured using reverberant shear wave elastography ex vivo [23], corresponding to SMR = *G*_*t*_*/G*_*p*_ *≈*2.49, further supported by neural network-based MRE estimates reporting a shear anisotropy parameter *ϕ* = *µ*_*L*_*/µ*_*T*_ – 1 = 1.09*±* 0.05 at 400 Hz [24], equivalent to SMR = *µ*_*L*_*/µ*_*T*_ *≈*2.09, collectively confirming that chicken breast SMR falls within the range of 2–3 and making it a well-characterized anisotropic tissue model for experimental validation of APRIL. ARFI excitations were applied at focal depth 25 mm across multiple angular orientations to identify the maximum and minimum peak displacement directions, from which the PDR was computed consistent with the acquisition protocol described above.

#### 2) Tissue-Mimicking Gelatin Phantom Fabrication

Two gelatin-based phantoms were fabricated for controlled validation. The background matrix for both phantoms was prepared to a target stiffness of ~10 kPa using: 41.273 g porcine gelatin, 62.749 g graphite powder (acoustic scatterers), 22.405 mL n-propanol (speed of sound correction), and 807.137 mL deionized water. Ingredients were dissolved at 60°C, cooled to ~37°C, and poured into molds.

##### i. Chicken Breast Gelatin Phantom

A trimmed chicken breast specimen was embedded as an anisotropic inclusion (SMR*≈*2–3) (section II-D-1) at a depth of 25 mm within the near-isotropic 10 kPa gelatin background (SMR *≈*1), with its dominant fiber axis aligned to a known laboratory reference direction.

##### ii. Isotropic Gelatin Inclusion Phantom (Negative Control)

A stiffer 40 kPa isotropic gelatin inclusion was fabricated separately using: 80.178 g porcine gelatin, 62.749 g graphite powder, 22.405 mL n-propanol, and 768.232 mL deionized water. The solidified inclusion was embedded within the 10 kPa background matrix. Since both components are isotropic gelatin with no fiber architecture, this phantom validated that APRIL does not artificially introduce anisotropy at mechanical discontinuities in isotropic media. All phantoms were imaged within 48 hours of fabrication at focal depth 25 mm using the same acquisition protocol described above.

#### 3) In Vivo Murine Tumor Model

A syngeneic 4T1 triple-negative breast cancer mouse model was used for in vivo validation. A suspension of 1 *×* 10^5^ 4T1 cells in 50 *µ*L phosphate-buffered saline (PBS) was injected subcutaneously into the right mammary fat pad of female BALB/c mice. ARFI imaging was performed at Day 5 to Day 28 post implantation in the same animal to capture the temporal evolution of tumor mechanical anisotropy. For imaging, the mouse was placed in a supine position under isoflurane anesthesia (3–5% induction, 1.5–2% maintenance) delivered via nose cone. The tumor-bearing region was depilated and ultrasound coupling gel was applied. ARFI excitations were delivered at a focal depth of 18 mm. All animal procedures were conducted in accordance with IACUC protocols.

Table I summarizes the complete training and test configurations used in this study.

**TABLE I.** Training and Test Dataset Summary.

| Parameter | Training (Homogeneous) | Test (Homogeneous) | Acoustic Variation Test | Test (Heterogeneous) | Experimental Validation (Biological Tissue) |
| --- | --- | --- | --- | --- | --- |
| Shear Moduli Ratio (SMR) | 0.9:0.25:4.9 | 1.25:0.5:4.25 | 1.75, 2.75, 3.75 | Inclusion: 1, 2, 4<br>Background: 1, 4 | Chicken breast: 2-3<br>Murine tumor: Unknown |
| Speed of Sound (m/s) | 1540 | 1540 | 1500, 1540, 1580 | 1540 | ~1540 |
| Attenuation (dB/cm/MHz) | 0.5 | 0.5 | 0.25, 0.5, 1.0 | 0.5 | ~0.5 |
| Density (kg/m<sup>3</sup>) | 1000 | 1000 | 900, 1000, 1100 | 1000 | ~1000 |
| Focal Depth (mm) | 18:2:30 | 22 | 22 | 25 | 18, 25 |

### E. Data Processing

Raw RF data were processed offline in MATLAB R2022b (MathWorks, Natick, MA). Axial displacement estimation was performed using one-dimensional normalized cross-correlation with sub-sample interpolation. Displacement waveforms were obtained as functions of axial depth and time.

For each depth, the peak displacement (PD) was identified from the displacement-time profile. The peak displacement ratio (PDR) was computed as the ratio of PD measured with the excitation beam PSF aligned parallel versus perpendicular to the axis of symmetry.

The level of asymmetry (LoA) was computed from simulated PSF beamwidths at corresponding depths. At inference, measured PDR profiles were matched to the precomputed LoA-conditioned PDR–SMR mappings generated during training. Depth-wise SMR values were then estimated and aggregated to form two-dimensional anisotropy maps. For experimental acquisitions, the LoA at each depth was estimated from the theoretical PSF beamwidths of the L11-5v transducer computed using Field II simulation under the acquisition parameters used (push frequency 4.75 MHz, F-number 1.5, focal depth 25 mm for phantoms and chicken breast; focal depth 18 mm for murine tumor). The simulated elevational and lateral −6 dB beamwidths were computed at each axial depth and used to derive the LoA profile, which was then applied to select the appropriate LoA-conditioned PDR–SMR mapping at inference. This approach is consistent with prior ARFI-based studies that employ simulated PSF characterization to support experimental displacement analysis [15], [16].

### F. Evaluation Metrics

Quantitative performance was assessed using mean percentage error (MPE), mean absolute error (MAE), root mean squared error (RMSE), and structural similarity index measurement (SSIM).

For homogeneous materials, predicted SMR profiles were compared against ground-truth SMR values across the axial depth range. For heterogeneous inclusion phantoms, evaluation was performed both globally and within manually defined inclusion regions of interest (ROIs).

SSIM was used to assess spatial fidelity of reconstructed anisotropy maps, while MPE quantified relative stiffness estimation error.

To evaluate robustness to acoustic variability, APRIL was tested under variations in speed of sound (1500–1580 m/s), attenuation (0.25–1 dB/cm/MHz), and density (900–1100 kg/m^3^). Performance metrics were computed under each condition to assess stability of the LoA-conditioned mapping.

## III. Result

### A. Depth-Dependent Behavior of PDR and LoA

The relationship between PDR and SMR was found to vary across depth, governed by the level of asymmetry (LoA) of the ARFI excitation PSF. Fig. 2 (top panel) shows LoA and PDR versus depth at a focal depth of 22 mm. The figure shows that LoA gradually increases toward the focus, reaching a peak where the elevational beamwidth is dominant, and then decreases beyond the focal zone. The corresponding PDR curve follows this trend, with relatively smooth variation near the focus but clear inflection points as depth moves into regions where the PSF geometry transitions (marked in the figure). Therefore, PDR = 1 when LoA = 1, PDR *>* 1 when LoA *>* 1, and vice versa. These findings justify the use of different regression models depending on LoA and depth.

Fig. 2 (bottom panel) presents PDR versus SMR curves at focal depths of 26 mm. Both data points and fitted models are shown, with polynomial regression yielding smooth mappings in stable LoA regions. The curve at depth 23 for the same focal depth, focuses on the transition regions (22-23.5 mm, and 29.5-31 mm for focal depth 26 mm) highlighting where spline interpolation provided a more accurate fit than polynomials. These deviations highlight the increased modeling challenge at depths where the PSF geometry alternates, as reflected in the figure by divergence between the data points and fitted curve. Both polynomial regression and spline interpolation are illustrated to understand how the polynomial regression fails and spline interpolation takes over.

### B. Depth-Wise Prediction Accuracy

Fig 3 shows the plot of predicted versus true SMR, along with line of equity (LoE) for focus at 22 mm. At the focal zone (19-22 mm), predictions clustered tightly around the LoE, with root mean squared errors (RMSE) lower than 0.1. At the transition region outside the focal depth (26 mm), error margins widened slightly with RMSE 0.13; reflecting the influence of PSF transitions, but predictions remained unbiased.

**Fig. 3.**
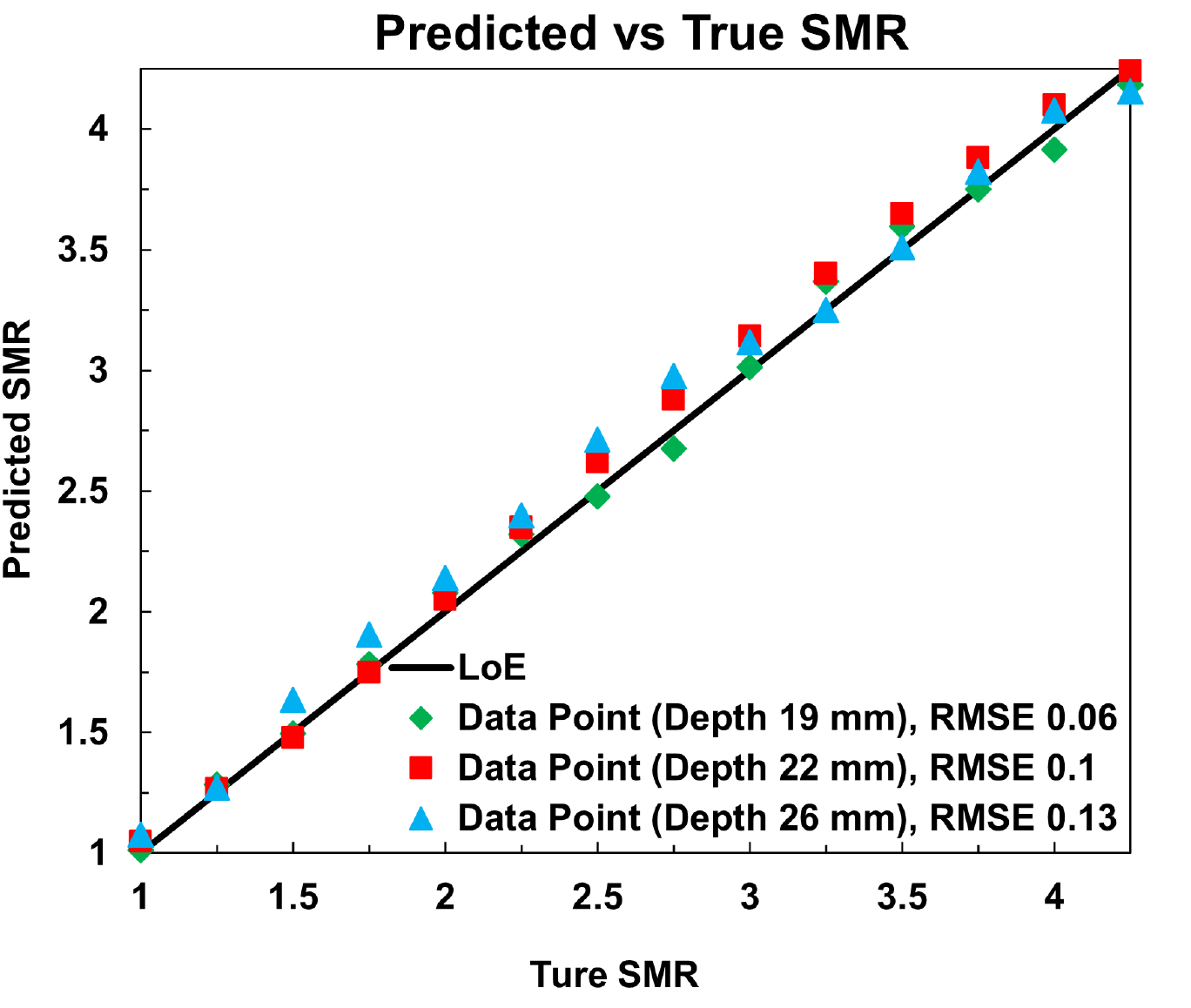
Predicted SMR vs True SMR for the test set at focal depth 22 mm, and two transition regions (19 mm & 26 mm) at both side of the focal depth along with line of equity (LoE).

To quantify prediction accuracy across depths, APRIL estimates were compared with ground truth SMR values at test depths of 10-30 mm for focus at 22 mm (Fig 4), with speed of sound 1540 m/s^2^, attenuation coefficient 0.5 dB/cm/MHz, and density 1000 kg/m^3^. The prediction error is less than 9% across the whole imaging depth. The error is lower (less than 8%) near the focal region (14-22 mm), whereas higher (less than 13%) outside of the focal region; comparatively higher at the denser region (24-30 mm). The mean absolute error across the whole region is 3% with a mean absolute error near the focal region is 2.3%.

**Fig. 4.**
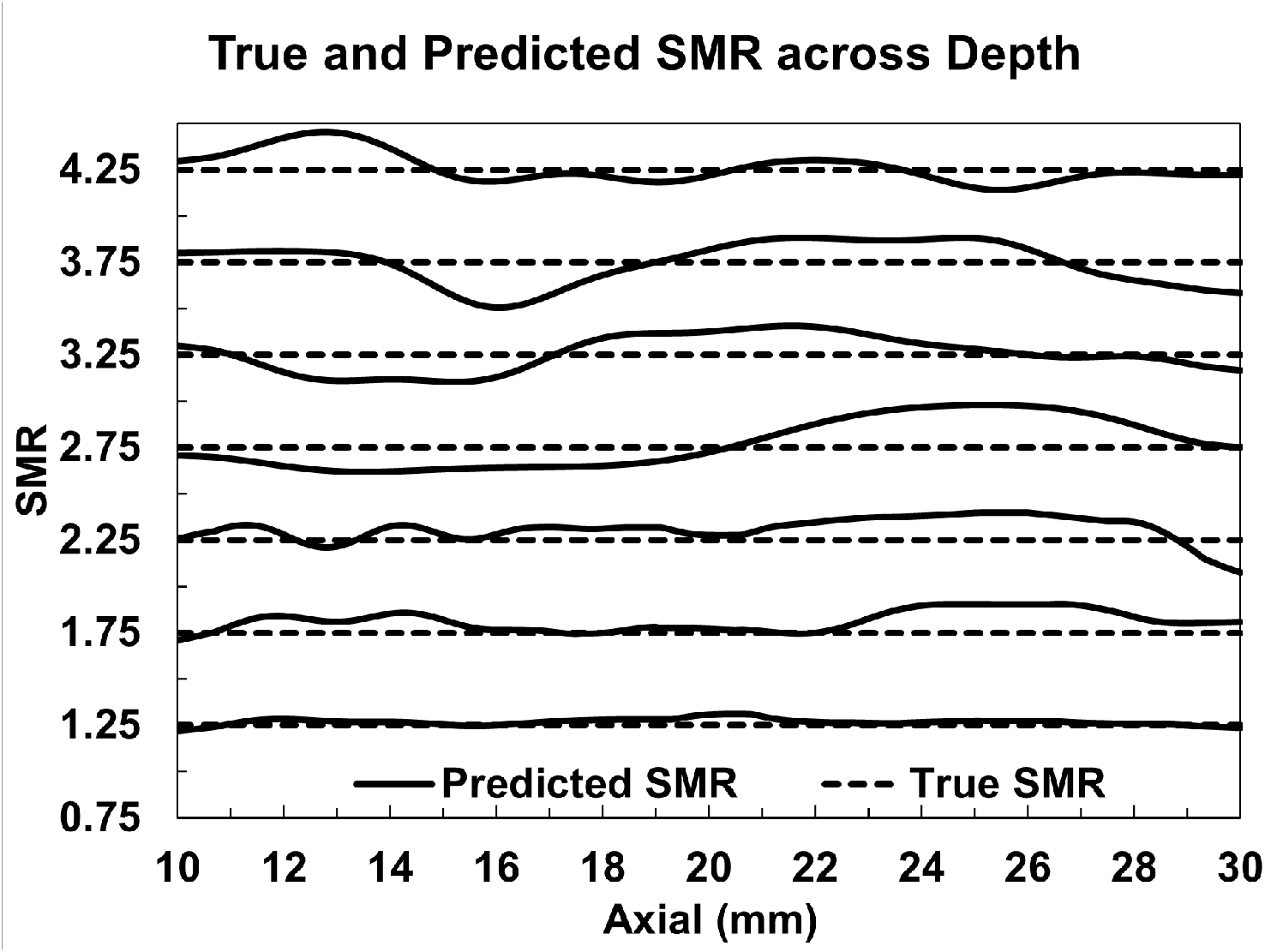
True SMR (TSMR) & predicted SMR (PSMR) across depth from 10 mm to 30 mm for focus 22 mm, SMR 1.25 to 4.25 with variation of 0.5, speed of sound 1540 m/s^**2**^, attenuation coefficient 0.5 dB/cm/MHz, and density 1000 kg/m^**3**^.

### C. Robustness to Acoustic Parameter Variations

Fig. 5 presents the predicted SMR distributions (Mean ± SD) under variations in speed of sound (1500, 1540, 1580 m/s), attenuation (0.25, 0.5, 1 dB/cm/MHz), and density (900, 1000, 1100 kg/m^3^) for true SMR values of 1.75, 2.75, and 3.75, evaluated over an axial depth range of 10–30 mm at focal depth 22 mm. Across all three acoustic parameter variations, APRIL consistently predicted SMR values close to their ground truth counterparts. For speed of sound variation (Fig. 5 (a)), predicted SMR distributions remained tightly grouped across all three conditions, with minor spread observed at higher true SMR (3.75), where the interquartile ranges slightly widened. Similarly, attenuation variation (Fig. 5 (b)) produced comparable median predictions across all conditions, though the highest attenuation (1 dB/cm/MHz) introduced marginally increased variance, particularly at true SMR = 3.75. Density variation (Fig. 5 (c)) exhibited a similar trend, with predictions remaining stable at lower SMR values while showing slightly elevated spread at true SMR = 3.75, notably for 1100 kg/m^3^. Across all conditions and parameter combinations, mean absolute errors remained below 10%, with the lowest variability observed at true SMR = 1.75 and the largest spread at true SMR = 3.75. It is worth noting that among all tested conditions, only the nominal acoustic parameter combination (1540 m/s, 0.5 dB/cm/MHz, 1000 kg/m^3^) was present in the training dataset, and none of the evaluated SMR values (1.75, 2.75, 3.75) were seen during training, making all reported results strictly out-of-distribution evaluations.

**Fig. 5.**
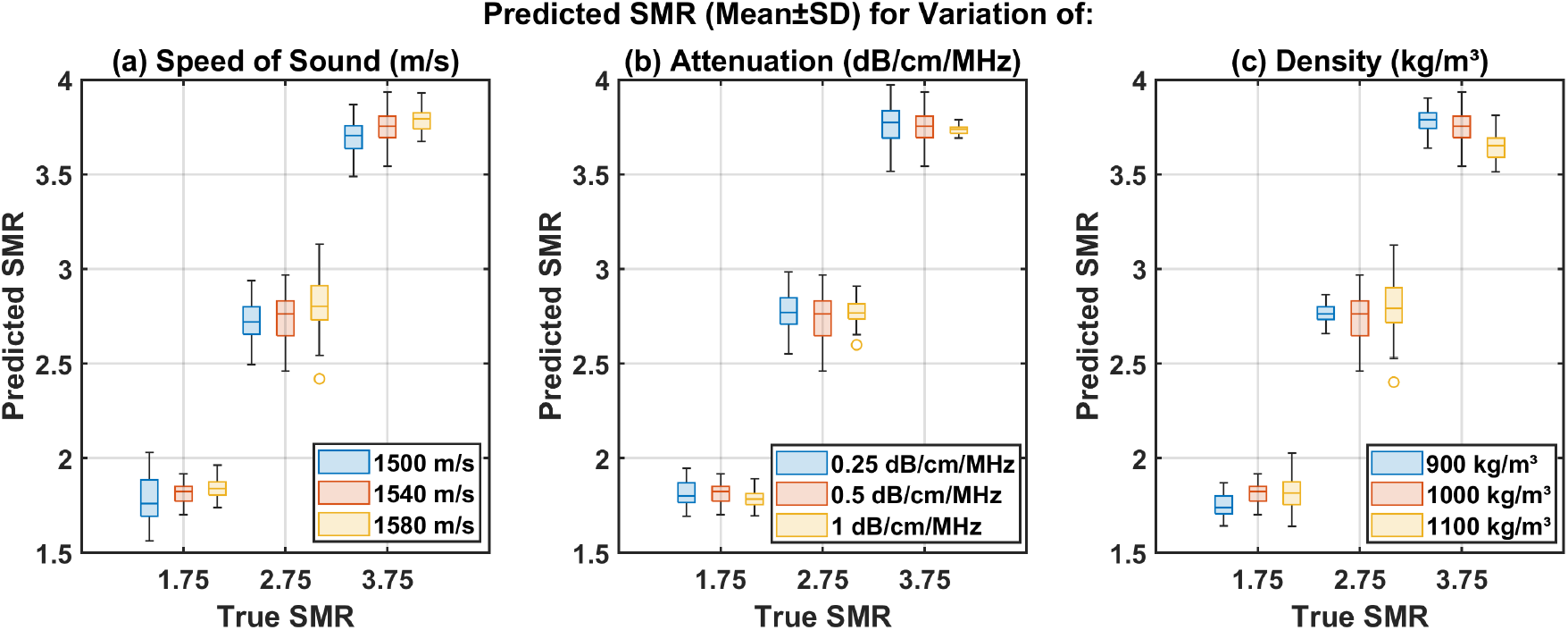
Predicted SMR (Mean ± SD) under variations in acoustic parameters over axial depth 10–30 mm at focal depth 22 mm for true SMR values of 1.75, 2.75, and 3.75: (a) speed of sound (1500, 1540, 1580 m/s), (b) attenuation coefficient (0.25, 0.5, 1 dB/cm/MHz), and (c) density (900, 1000, 1100 kg/m^3^). Note that only the combination of 1540 m/s, 0.5 dB/cm/MHz, and 1000 kg/m^3^ was included in the training dataset but none of the evaluated SMR values (1.75, 2.75, 3.75) were present during training, confirming that all test conditions represent out-of-distribution generalization.

### D. Inclusion Case Reconstruction

Fig. 6 presents reconstructed two-dimensional (2D) anisotropy maps for three heterogeneous phantoms containing inclusions with different shear-modulus ratios (SMR) embedded within isotropic backgrounds, and vice versa. Each map corresponds to a unique inclusion-background (I-B) stiffness contrast: (*I* = 4, *B* = 1), (*I* = 2, *B* = 1), and (*I* = 1, *B* = 4), where 1 indicates isotropic and higher than 1 indicates anisotropic. The inclusions were centered at a focal depth of 25 mm with a radius of 3 mm.

**Fig. 6.**
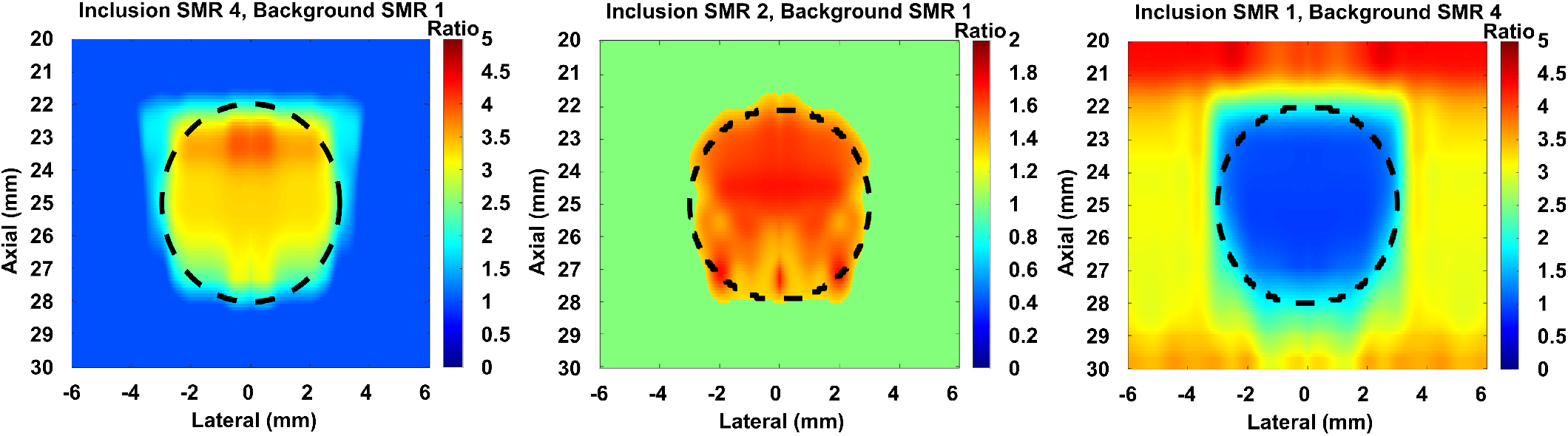
Reconstructed 2D SMR maps for three heterogeneous phantom configurations centered at focal depth 25 mm with inclusion radius 3 mm: (left) anisotropic inclusion (SMR = 4) in isotropic background (SMR = 1); (center) moderate-contrast anisotropic inclusion (SMR = 2) in isotropic background (SMR = 1); (right) isotropic inclusion (SMR = 1) in anisotropic background (SMR = 4). Dashed contours indicate the true inclusion boundaries. Color bars indicate predicted SMR values.

In the high-contrast case (*I* = 4, *B* = 1), the inclusion boundary was sharply defined and the reconstructed SMR distribution closely matched the ground truth with SSIM = 83% and an overall MPE = 8.72%. Region-specific analysis revealed that the reconstruction error was primarily concentrated within the inclusion (inclusion MPE *≈*12.5%, estimated average SMR *≈*3.5), while the background region was recovered with high accuracy (background MPE *≈*7%, estimated average SMR *≈*1.07), with residual error attributable to boundary transition artifacts. For the moderate-contrast case (*I* = 2, *B* = 1), APRIL preserved both the inclusion geometry and magnitude with improved fidelity (SSIM = 86.1%, overall MPE = 6.44%). The inclusion region showed moderate underestimation (inclusion MPE *≈*10%, estimated average SMR *≈*1.8), whereas the isotropic background was reconstructed with notably high accuracy (background MPE *≈*5%, estimated average SMR *≈*1.05), reflecting the model’s stability at lower anisotropy contrasts.

When the inclusion was isotropic and embedded in a stiffer anisotropic background (*I* = 1, *B* = 4), the reconstruction dynamics were reversed. The isotropic inclusion was captured with excellent precision (inclusion MPE *≈*2%, estimated average SMR *≈*1.02), while the anisotropic background exhibited relatively higher reconstruction error (background MPE *≈*10%, estimated average SMR *≈*3.6), consistent with the overall MPE of 8.66% and the cross-boundary smoothing effect noted previously.

Overall, the figure demonstrates that APRIL effectively reconstructs anisotropic inclusions within isotropic and anisotropic backgrounds. Notably, the lower-SMR region whether inclusion or background is consistently reconstructed with greater accuracy (~ 2–7% MPE), while the higher-SMR region carries the dominant reconstruction error (~ 10–12.5% MPE). These results highlight the framework’s ability to generalize from homogeneous training data to complex heterogeneous tissue structures, while also revealing a systematic trend of higher error in regions with elevated anisotropy.

### E. Experimental Validation

Fig. 7 and 8 presents the 2D SMR maps overlaid on B-mode images acquired experimentally using an L11-5v transducer across five specimens: ex-vivo chicken breast, a tissue mimicking gelatin isotropic phantom, chicken breast embedded in gelatin phantom, and in-vivo murine tumor at Day 5 and Day 28 post 4T1 cell implantation. In the murine tumor model, APRIL produced spatially coherent SMR maps that clearly differentiated the tumor boundary (dashed contour) from the surrounding tissue on both imaging days. At Day 5, the tumor interior exhibited elevated SMR values predominantly in the range of 2.5-3, indicating a highly anisotropic mechanical environment consistent with early-stage tumor microstructural organization and collagen fiber alignment. At Day 28, the tumor had visibly grown in both axial and lateral extent, and the reconstructed SMR map reflected a broader high-anisotropy region (SMR 3.5-4.5) with a more spatially expansive core, suggesting progressive microstructural remodeling associated with tumor proliferation. The surrounding tissue outside the dashed contour showed consistently lower SMR values (1-2), providing clear contrast between tumor and background. Critically, the evolution in SMR spatial distribution between Day 5 and Day 28 in the same animal demonstrates APRIL’s sensitivity to longitudinal changes in tissue anisotropy, highlighting its potential as a biomarker for tumor progression monitoring. In the ex-vivo chicken breast specimen, the reconstructed SMR map revealed spatially stratified anisotropy patterns aligned with the dominant muscle fiber orientation, with SMR values ranging from approximately 2-3, supported by the previously reported SMR in chicken breast [23], [24]. The layered structure of the SMR distribution visually corresponded to the fiber bundle architecture visible in the B-mode image, confirming APRIL’s ability to resolve directional mechanical heterogeneity within fibrous tissue. In the chicken breast gelatin phantom, APRIL reconstructed a localized high-anisotropy region (SMR 2-3) within the dashed ROI corresponding to the embedded chicken breast portion, while the surrounding gelatin background exhibited substantially lower SMR values (around 1) near isotropic. The spatial localization of the anisotropic inclusion within the isotropic gelatin matrix was clearly captured, demonstrating the method’s applicability to tissue-mimicking phantom configurations. In the isotropic gelatin phantom, the reconstructed SMR map showed uniformly low values (1-1.1) throughout the imaging field, with the dashed ROI confirming near-unity SMR within the isotropic inclusion-consistent with the isotropic mechanical nature of isotropic tissue. This result serves as a negative control, validating that APRIL does not artificially introduce anisotropy in mechanically isotropic specimens.

**Fig. 7.**
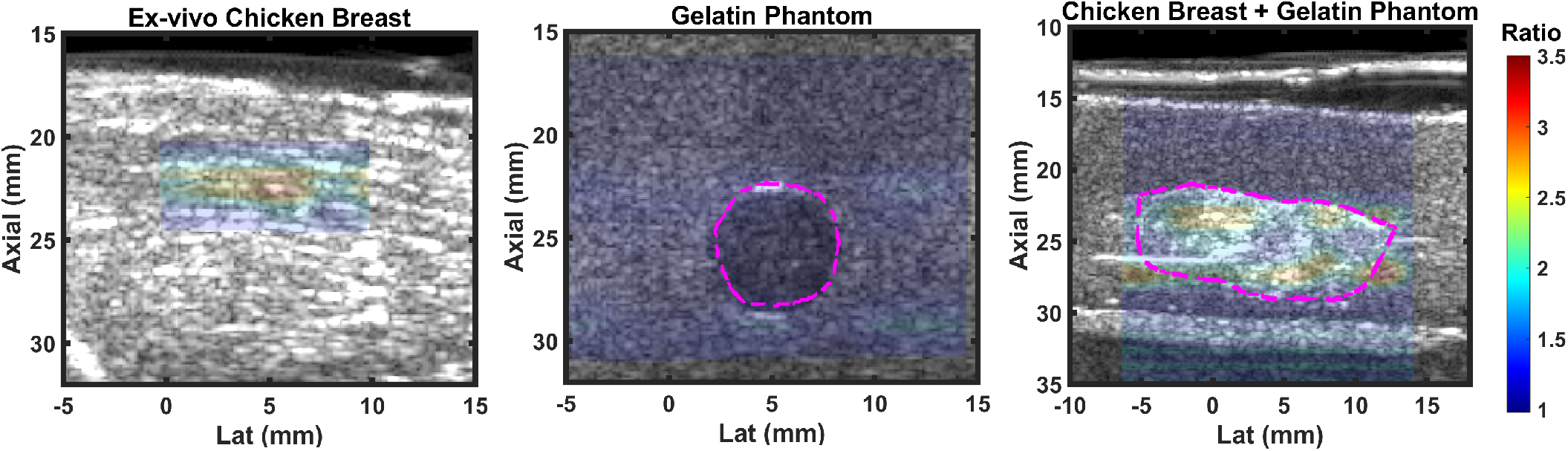
APRIL-predicted 2D SMR maps overlaid on B-mode images for ex vivo and phantom specimens acquired using an L11-5v transducer. (Left) Ex vivo chicken breast showing depth-stratified anisotropy aligned with dominant muscle fiber orientation (SMR 2-3). (Center) Isotropic gelatin inclusion phantom (40 kPa inclusion in 10 kPa background) confirming near-unity SMR (1-1.1) throughout, serving as a negative control. (Right) Chicken breast embedded in gelatin phantom (focal depth 25 mm) showing localized anisotropic inclusion (SMR 2-3) within near-isotropic gelatin background. For all specimens, PDR was computed from maximum and minimum peak displacements identified across multiple ARFI excitation orientations.

**Fig. 8.**
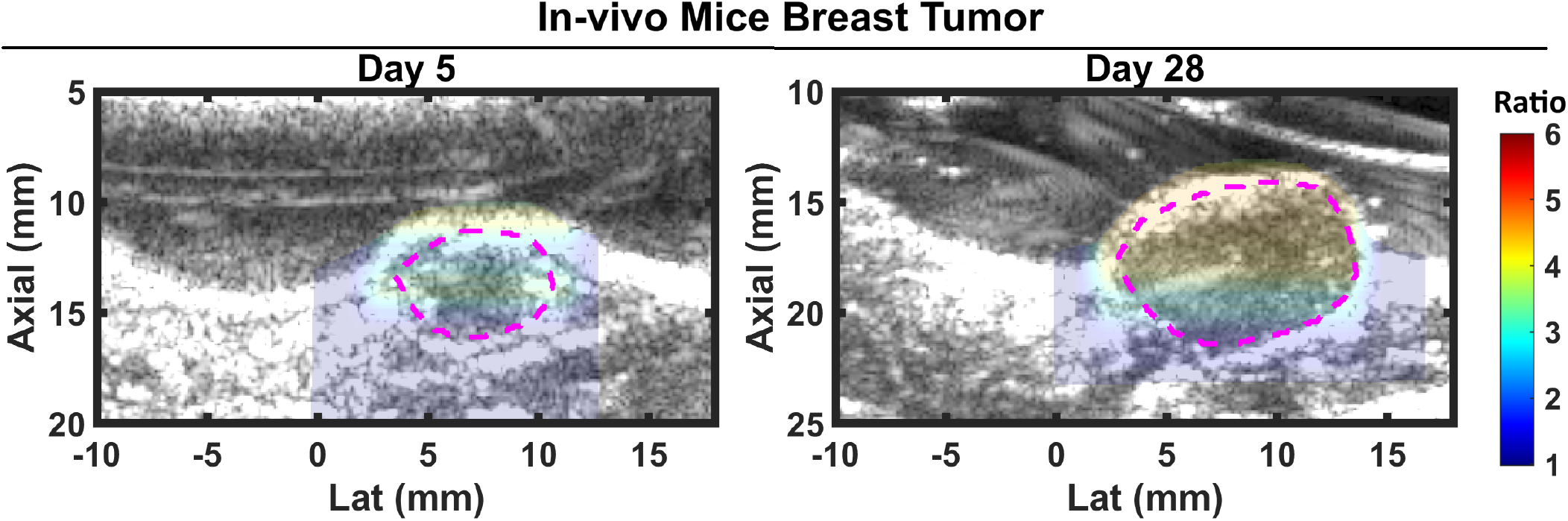
APRIL-predicted 2D SMR maps overlaid on B-mode images for in vivo murine 4T1 tumor model acquired using an L11-5v transducer at focal depth 18 mm. (Left) Day 5 post implantation showing early-stage tumor boundary (dashed contour) with near-isotropic SMR (2.5-3) distribution. (Right) Day 28 post implantation in the same animal showing progressive SMR elevation (3.5-4.5) and spatial expansion of the high-anisotropy region within the tumor interior, consistent with extracellular matrix remodeling and collagen fiber alignment associated with 4T1 tumor proliferation. PDR was computed from maximum and minimum peak displacements identified across multiple ARFI excitation orientations relative to the tumor boundary.

## IV. Discussion

This study presented APRIL, an adaptive regression-based framework for quantitative and depth-resolved anisotropy mapping using acoustic radiation force impulse (ARFI) displacements. By integrating polynomial regression and shape-preserving spline interpolation, APRIL establishes a continuous mapping between the peak displacement ratio (PDR) and the shear modulus ratio (SMR) across depth. The framework successfully extended prior focal-only anisotropy estimation methods into full two-dimensional (2D) imaging, demonstrated robustness under acoustic heterogeneity, and showed feasibility in both simulated and in vivo settings.

Conventional ARFI- and SWE-based approaches quantify anisotropy through point or focal-depth measures, often assuming a uniform polynomial relation between PDR and SMR. In contrast, APRIL adaptively identifies the appropriate regression type—polynomial or spline based on the level of asymmetry (LoA) of the excitation beam, which varies with depth due to geometric focusing. This LoA-conditioned modeling represents a key advance, enabling accurate reconstruction of SMR where prior empirical models fail due to PSF asymmetry or non-monotonic displacement behavior. The approach does not require training with heterogeneous media, yet it generalizes effectively to inclusion and in-vivo/ex-vivo cases, underscoring its physical interpretability and data efficiency.

In homogeneous materials, APRIL produced near-ideal SMR reconstructions (MPE *<* 2.3% near focal region), confirming that the adaptive polynomial-spline mechanism reliably captures the depth-varying PDR–SMR trend. Across multiple focal depths, errors remained lowest in the focal zone, consistent with the region of maximal acoustic energy and well-defined LoA. Slightly higher errors at off-focus depths (RMSE = 0.13) likely reflect reduced signal-to-noise ratio and the transition in PSF geometry.

The robustness of APRIL under acoustic parameter variations underscores a key advantage of the LoA-conditioned regression framework. Unlike purely data-driven approaches that may overfit to specific acoustic conditions present in training data, APRIL’s adaptive selection between polynomial regression and PCHIP spline interpolation provides a physically grounded mapping that is inherently less sensitive to moderate perturbations in tissue acoustic properties. The slightly elevated prediction variance observed at higher SMR values (3.75) across all three parameter conditions is consistent with the broader dynamic range of displacement responses at greater anisotropy levels, where small changes in acoustic parameters can produce proportionally larger perturbations in the PDR profile. The increased spread at higher attenuation (1 dB/cm/MHz) is expected, as greater signal attenuation reduces effective SNR at deeper imaging depths, thereby increasing uncertainty in peak displacement estimation. Similarly, the minor variation introduced by density and speed of sound changes is attributable to their influence on the acoustic radiation force distribution and the resulting tissue displacement magnitude, both of which modulate the PDR without fundamentally altering its monotonic relationship with SMR. Importantly, since APRIL’s training data were generated under fixed nominal acoustic conditions, these results demonstrate generalization beyond the training distribution, highlighting the framework’s suitability for clinical scenarios where precise knowledge of tissue acoustic properties is unavailable. When tested under acoustic parameter variations (speed of sound, attenuation, and density), APRIL maintained stable SMR profiles with mean absolute errors below 10%, demonstrating robustness to system and tissue heterogeneity. The use of LoA conditioning and shape-preserving interpolation prevented model drift under acoustic perturbations-a limitation often reported in purely empirical or deep learning–based elastography models.

The inclusion reconstruction results demonstrate that despite being trained exclusively on homogeneous media, APRIL generalizes effectively to heterogeneous tissue configurations—a practically significant outcome given that real biological tissues frequently contain localized anisotropic regions. The superior performance in the moderate-contrast case (*I* = 2, *B* = 1) compared to the high-contrast case (*I* = 4, *B* = 1) can be attributed to the fact that both inclusion and background SMR values fall within the well-supported interior of the spline training range, where the PDR-SMR mapping is most reliable. In contrast, the high-contrast case involves SMR = 4, which approaches the upper boundary of the training distribution (SMR up to 4.9), where spline extrapolation uncertainty is inherently higher. The lowest performance in the reversed-contrast case (*I* = 1, *B* = 4) can be explained by the nature of the inclusion–background interface. When an isotropic inclusion is surrounded by a stiffer anisotropic background, the physical signal difference between the two regions at the boundary is relatively small, making it harder for APRIL to distinguish one region from the other. This causes the reconstructed values near the boundary to blend across it, producing the smoothing artifacts visible in the figure (Fig. 6 (Right)).

Region-specific analysis further reveals a consistent systematic trend: the lower-SMR region whether inclusion or background-is reconstructed with greater accuracy (~7% MPE), while the higher-SMR region carries the dominant reconstruction error (~12.5% MPE). This asymmetry suggests that APRIL’s PDR-SMR mapping has inherently higher sensitivity in resolving lower anisotropy values, likely due to denser sampling of the training distribution in that range. Importantly, in the reversed-contrast case (*I* = 1, *B* = 4)-which most closely mimics clinically relevant scenarios such as isotropic tumors embedded within anisotropic healthy tissue-the inclusion itself was reconstructed with the highest regional accuracy of all three configurations (inclusion MPE *≈* 2%), despite the lower overall SSIM. This indicates that APRIL prioritizes accurate delineation of the pathological region of interest even when background reconstruction fidelity is reduced, which is of greater diagnostic relevance in practical imaging scenarios. Nevertheless, SSIM values ranging from 75.3% to 86.1% and MPE below 9% across all configurations confirm that APRIL maintains clinically meaningful spatial fidelity and contrast sensitivity, underscoring its potential for imaging heterogeneous tissue pathologies such as fibrotic inclusions, focal muscle lesions, and tumor boundaries. The systematic underestimation observed in high-SMR regions further suggests that augmenting the training dataset with additional high-anisotropy samples near the upper boundary of the SMR range could further improve reconstruction fidelity in extreme-contrast configurations.

The experimental results collectively validate APRIL across a diverse range of tissue types and mechanical configurations, spanning anisotropic fibrous tissue, tumor-bearing tissue, tissue-mimicking phantoms, and isotropic controls-each presenting distinct imaging challenges that a robust anisotropy mapping framework must address. The murine tumor data represent the most significant experimental finding of this study. The ability of APRIL to detect and spatially map elevated anisotropy within the tumor interior, and to track the longitudinal evolution of that anisotropy between Day 5 and Day 28 in the same animal, demonstrates a capability that is not achievable with conventional focal or point-wise ARFI and SWE methods. The elevated SMR observed within the tumor is consistent with the known biomechanics of 4T1 breast tumors, which are characterized by progressive extracellular matrix remodeling, increased collagen deposition, and fiber alignment-all of which contribute to directional stiffness anisotropy. The expansion of the high-SMR region between Day 5 and Day 28 parallels the physical growth of the tumor visible in the B-mode images, suggesting that the SMR maps reflect genuine microstructural progression rather than imaging artifacts. This finding positions APRIL as a potentially valuable non-invasive tool for longitudinal tumor characterization, complementary to conventional stiffness-based elastography by adding a directional mechanical dimension that may more sensitively reflect early-stage pathological remodeling. The chicken breast results further reinforce the method’s biological relevance. The layered SMR distribution observed in the ex-vivo specimen reflects the known hierarchical organization of skeletal muscle-comprising fascicles, fiber bundles, and individual fibers-each contributing to the depth-dependent anisotropy pattern. The coherence between the SMR map and the B-mode structural appearance confirms that APRIL resolves biologically meaningful mechanical gradients rather than PSF or noise artifacts. The phantom experiments provide controlled validation of APRIL’s contrast sensitivity and specificity. The chicken breast gelatin phantom confirmed that APRIL can localize an anisotropic inclusion within an isotropic background in a controlled setting, consistent with the simulation-based inclusion results in Section III-D. The isotropic gelatin phantom, serving as an isotropic negative control, is particularly important: the near-unity SMR values throughout the field confirm that APRIL does not introduce spurious anisotropy estimates in mechanically isotropic media, which is a critical requirement for clinical credibility. Together, these experimental results spanning simulated, phantom, ex-vivo, and in-vivo conditions-establish APRIL as a generalizable and physically interpretable framework for quantitative 2D anisotropy imaging with direct translational relevance to musculoskeletal, oncological, and organ biomechanics applications.

Two practical limitations of the current APRIL framework warrant discussion. First, accurate SMR estimation requires ARFI excitations applied precisely parallel (0°) and perpendicular (90°) to the tissue axis of symmetry (AoS)-orientations that are not always identifiable a priori in clinical settings. In the present study, this limitation was addressed experimentally by acquiring ARFI displacement responses across multiple angular orientations relative to the presumed dominant fiber direction, and computing the peak displacement ratio as the ratio of maximum to minimum peak displacement across all acquired angles. While this multi-angle acquisition strategy provides a practical approximation of the true parallel and perpendicular responses, it cannot guarantee exact 0° and 90° alignment, and the resulting PDR may therefore represent a close but not exact estimate of the true SMR-linked displacement ratio. Future work will focus on training a model to learn the full peak displacement versus angle pattern from multi-orientation acquisitions, enabling automatic identification of the true maximum and minimum peak displacement directions without requiring prior knowledge of the tissue AoS-thereby eliminating this orientation dependency entirely. Second, the current framework requires prior knowledge of the LoA profile across depth for the specific transducer model and acoustic parameter configuration being used. In this study, LoA was derived from Field II simulations of the L11-5v transducer under the specific acquisition parameters employed. For a different transducer model, focal configuration, or substantially different tissue acoustic properties, a new set of LoA-conditioned PDR–SMR mappings would need to be simulated and stored, limiting the out-of-the-box transferability of the trained model to new imaging setups. To overcome this constraint, future work will explore the development of a generalized deep learning framework that jointly learns the relationship between PSF geometry, acoustic parameters, and PDR-SMR mappings across a broad range of transducer configurations-eliminating the need for setup-specific simulation pipelines and enabling direct deployment across diverse clinical ultrasound systems without retraining.

## V. Conclusion

This study presented APRIL, a quantitative, depth-resolved 2D anisotropy imaging framework that extends ARFI-based focal degree-of-anisotropy estimation into full two-dimensional shear modulus ratio mapping. By adaptively selecting between polynomial regression and shape-preserving spline interpolation conditioned on the level of asymmetry (LoA) of the ARFI excitation PSF, APRIL addresses the fundamental challenge of depth-varying PDR–SMR relationships that arise from geometric beam focusing-a limitation that prior focal and point-wise anisotropy estimation methods did not overcome. Across simulation-based evaluations, APRIL achieved depth-resolved SMR prediction errors below 9% over an axial range of 10–30 mm, with highest accuracy in the focal region (MAE 2.3%, RMSE *<* 0.1). In heterogeneous inclusion phantoms, APRIL reconstructed spatially coherent anisotropy maps with SSIM up to 86.1% and MPE below 7%, accurately delineating inclusion boundaries despite being trained exclusively on homogeneous media-demonstrating strong generalization to unseen tissue configurations. Under acoustic parameter variations spanning speed of sound, attenuation, and density, mean absolute errors remained below 10%, confirming robustness to system and tissue heterogeneity that is essential for clinical deployment. Experimental validation across diverse specimen types-including ex vivo chicken breast, tissue-mimicking phantoms, an isotropic gelatin control, and in-vivo murine tumor model-further established the practical viability of APRIL. Most significantly, the longitudinal murine tumor experiment demonstrated APRIL’s sensitivity to progressive microstructural remodeling between Day 5 and Day 28 post tumor implantation, revealing spatially expanding high-anisotropy regions consistent with collagen fiber alignment and extracellular matrix remodeling associated with 4T1 tumor proliferation. This capability of tracking the spatial evolution of tissue anisotropy over time in a living subject-represents a meaningful advance over existing elastography methods that report only scalar stiffness or focal anisotropy estimates. Collectively, these results establish APRIL as a physically grounded, generalizable, and clinically viable framework for quantitative anisotropy biomarker imaging. Its applicability spans a broad range of tissues and pathologies-including skeletal muscle disease, tendon injury, renal fibrosis, myocardial remodeling, and tumor characterization — where directional mechanical properties carry diagnostic and prognostic significance. Future work will focus on extending APRIL to in vivo human studies, incorporating multi-focal zone acquisitions to further improve depth coverage, and integrating the framework with real-time ultrasound platforms to enable clinical translation.

## VI. Acknowledgment

This work was supported in part by a Cancer Center pilot grant funded through the NIH National Cancer Institute Cancer Center Support Grant P30CA071789, and by Dr. Hossain’s startup award from the University of Hawaii at Mānoa College of Engineering. The technical support and advanced computing resources from University of Hawaii Information Technology Services - Research Cyberinfrastructure, funded in part by the National Science Foundation CC* awards 2201428 and 2232862 are gratefully acknowledged.

